# The Hox Transcription Factor Ubx stabilizes Lineage Commitment by Suppressing Cellular Plasticity

**DOI:** 10.1101/437947

**Authors:** Katrin Domsch, Julie Carnesecchi, Vanessa Disela, Jana Friedrich, Nils Trost, Olga Ermakova, Maria Polychronidou, Ingrid Lohmann

## Abstract

During development cells become gradually restricted in their differentiation potential by the repression of alternative cell fates. While we know that the Polycomb complex plays a crucial role in this process, it still remains unclear how alternative fate genes are specifically targeted for silencing in different cell lineages. We address this question by studying Ultrabithorax (Ubx), a multi-lineage transcription factor (TF) of the Hox class, in the mesodermal and neuronal lineages using sorted nuclei of *Drosophila* embryos and by interfering with Ubx in mesodermal cells that have already initiated differentiation. We find that Ubx is a key regulator of lineage development, as its mesoderm-specific depletion leads to the de-repression of many genes normally expressed in other lineages. Ubx silences expression of alternative fate genes by interacting with and retaining the Polycomb Group (PcG) protein Pleiohomeotic (Pho) at Ubx targeted genomic regions, thereby setting repressive chromatin marks in a lineage-dependent manner. In sum, our study demonstrates that Ubx stabilizes lineage choice by suppressing the multi-potency encoded in the genome in a lineage-specific manner via its interaction with Pho. This mechanism may explain why the Hox code is maintained throughout the lifecycle, since it seems to set a block to transdifferentiation in many adult cells.

## INTRODUCTION

Multicellular animals owe their complexity to their capacity to produce and maintain different lineages composed of multiple cell types that share virtually the same genomic DNA. Such complexity requires a tight regulation of gene expression to unambiguously specify and constrain the developmental path taken by cells within and between different lineages. Cell fates are controlled by networks of transcription factors (TFs) that act in the context of the genomic chromatin state to activate transcriptional programs realizing the distinct properties of cells of a given lineage^1–3^. The forced expression of TFs expressed only in one lineage like Myoblast Determination Protein (MyoD), which is sufficient to alter cell identity^4,5^, showed that lineage-restricted TFs can act as major lineage switches by activating lineage-specific transcriptional programs. However, recent profiling experiments revealed that global gene expression and chromatin accessibility changes are imperfect even in MyoD reprogrammed cells, showing that these highly potent reprogramming TFs cannot completely erase the original cell or lineage fate and unequivocally induce the new one^6^. In addition, lineage choice does not only involve the activation of one lineage-specific transcriptional program but also the repression of the programs of all alternative lineages^7,8^, a regulatory wiring that needs to be faithfully accomplished in all the different lineages. Although lineage-restricted TFs could in principle fulfil this function, it would ask for a highly complex regulatory architecture. On the other hand, TFs expressed in multiple/all lineages with a rather promiscuous binding behaviour would dramatically reduce this complexity and would elegantly explain lineage-specific gene regulation based on the interaction of these broadly expressed TFs with lineage-restricted factors.

Hox TFs represent an excellent model to address this fundamental question, since they are active in all lineages along the anterior-posterior (AP) axis of bilaterian animals. Importantly, the famous and well-known identity switch of whole body parts, the homeotic transformation, which is induced by altered Hox expression^9,10^, shows that Hox TFs have comparable functions in all lineages during development and indicates that they control the development of different lineages in a highly specific manner. The latter conclusion is supported by recent experiments showing that mesoderm-derived vascular wall mesenchymal stem cells (VW-MSCs) can be generated *in vitro* from induced pluripotent stem cells (iPSCs) simply by inducing the expression of a mixture of *Hox* genes that are selectively expressed in adult VM-MSCs^11^. However, while this study showed that *Hox* genes alone are sufficient to induce the generation of one specific cell type of one lineage *in vitro*, it left the questions open whether this is also the case *in vivo*, and how Hox TFs unambiguously select among the many possible transcriptional programs only one, which drives a cell or a whole lineage into one specific direction.

One major bottleneck in this direction is that genome-wide Hox chromatin binding studies have been performed so far mainly in cell culture systems^12,13^, specialized epithelial tissues^14^, or mixtures of cell lineages^15^, hampering the identification of common and lineage-specific mechanisms employed by Hox TFs in different lineages *in vivo*. Furthermore, unlike lineage-restricted TFs, which are often tested *in vivo* using ectopic expression systems^16,17^, the functional analysis of TFs acting in multiple lineages requires the targeted interference with these factors in individual lineages. With the availability of conditional genome editing^18,19^ and nanobody driven protein degradation systems^20^ this is now possible in an efficient manner and allows to elucidate the mode of action of Hox TFs in individual lineages, which are located in an otherwise unperturbed tissue environment at any stage in development. This is particularly important for multi-lineage TFs like the Hox proteins, which extensively control cell communication^21,22^ and thus influence their own action in neighbouring lineages.

Here, we probe the Hox TF Ubx in the mesodermal and neuronal lineages using sorted nuclei of *Drosophila* embryos and by interfering with Ubx function specifically in the mesoderm lineage that is specified and fully committed to the mesodermal fate. To this end, we generated a GFP-Ubx gene fusion at the endogenous locus using CRISPR-Cas9 and homologous recombination. We show that Ubx is a key regulator of lineage development and diversification, as it controls the mesodermal and neuronal transcriptional programs with high specificity despite interacting with many genes in both lineages. Intriguingly, our study demonstrates that Ubx controls lineage specification and differentiation by restricting cellular plasticity within each lineage. In the mesoderm Ubx executes this function by silencing alternative fate genes, and we show that this repression requires the interaction of Ubx with the Polycomb Group (PcG) protein Pleiohomeotic (Pho) on co-bound chromatin sites. We furthermore find that Ubx stabilizes Pho binding to Ubx targeted genomic regions, and that this interaction is critical for gene repression by controlling the balance of H3K27me3 as well as H3K27ac at these sites. Taken together, our study demonstrates that Hox TFs control not only the segmental identity but also the developmental programs intrinsic to each lineage with high precision, and that one of the prominent functions of this multi-lineage TF family is the repression of alternative lineage programs, which restricts cellular plasticity in a lineage-specific manner.

## RESULTS

### A comprehensive map of transcriptional profiles and Ubx binding in the *Drosophila* embryonic mesodermal and neuronal lineages

To elucidate mechanisms that enable a multi-lineage TF to instruct the development of divergent lineages, we recorded on the tissue level chromatin binding of the broadly expressed Hox protein Ubx in the mesoderm and nervous system and determined the transcriptomes of these two lineages during two stages of *Drosophila* embryogenesis. To this end, we used the <u>i</u>solation of <u>n</u>uclei <u>ta</u>gged in specific <u>c</u>ell <u>t</u>ypes (INTACT) method^13^ by inducing the expression of the nuclear membrane protein Ran GTPase activating protein (RanGAP) fused to a biotin ligase acceptor peptide and the *E. coli* Biotin ligase (BirA) in the mesoderm via the pan-mesodermal *twist* (*twi*)^14^ or the pan-neuronal *embryonic lethal abnormal vision* (*elav*)^25,26^ regulatory regions. Co-expression of both transgenes in *Drosophila* embryos, which did not affect development (Supplementary Fig. 1C-1H), resulted in cell type-specific biotinylation of nuclei *in vivo* (Fig. 1A-C’). Mesodermal or neuronal nuclei were efficiently isolated using Streptavidin-coated beads, as we detected Mef2-positive nuclei and concomitantly mesodermal marker gene expression like *Mef2 and nautilus* (*nau*) almost exclusively in the mesodermal collection (Fig. 1D), while Elav-labelled neuronal nuclei and neuronal marker genes like *eagle* (*eg*) and *elav* were found highly enriched in the neuronal fraction only (Supplementary Fig. 1A-B).

**Fig. 1:**
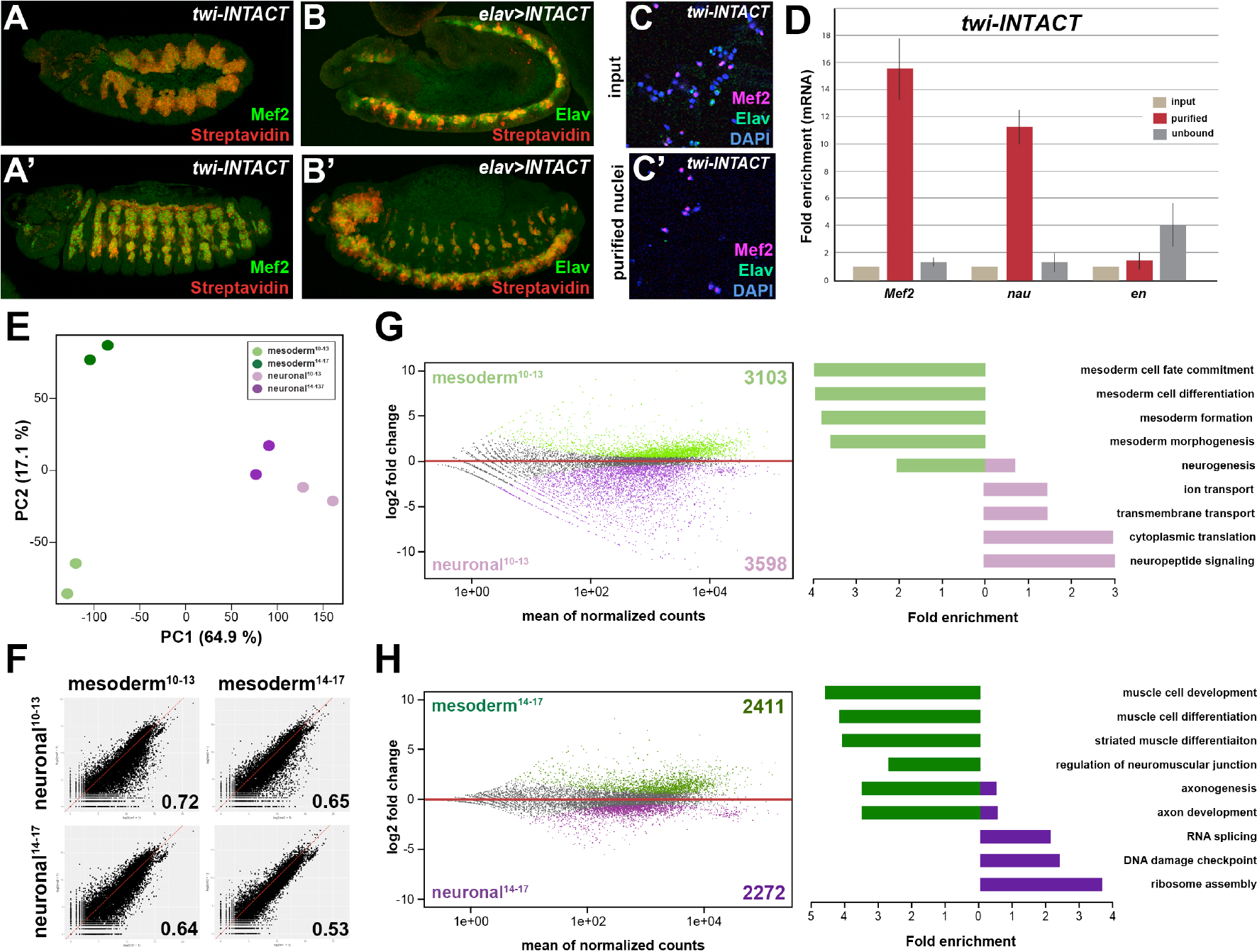
Comparative profiling of the mesodermal and neuronal transcriptomes in *Drosophila*. **embryos (A, A’)** Lateral views of stage 11 (A) and stage 14 (A’) *twi-INTACT Drosophila* embryos stained for the muscle differentiation marker Mef2 (green) and Streptavidin (red). **(B, B’)** Lateral view of stage 11 (B) and stage 14 (B’) *elav>INTACT Drosophila* embryos stained for the pan-neuronal marker Elav (green) and Streptavidin (red). **(C, C’)** Nuclei obtained from *twi-INTACT* embryos stained for Mef2 (red), Elav (green) and DAPI (blue) before (C) and after (C’) INTACT purification. **(D)** Expression of mesodermal (*Mef2*, *nau*) and ectodermal (*en*) mRNA transcripts in nuclei purified from *twi*-INTACT embryos as measured by RT-PCR using RNA from total/input (10%), purified and unbound nuclei. **(E)** Principle Component Analysis (PCA) applied to all RNA-seq samples identifies two clusters (mesodermal, neuronal) sharing similar expression signatures. **(F)** Pearson correlation analysis highlights the differences and similarities between the mesodermal and neuronal RNA-seq datasets. **(G, H)** Left: MA analysis identifies genes differentially expressed in the mesodermal and neuronal lineages at two different time windows, embryonic stage 10-13 (G) and embryonic stage 14-17 (H). Genes differentially expressed in the mesodermal lineage are indicated as green dots, those in the neuronal lineage as purple dots. Right: Bar diagram displaying fold enrichment of gene ontology terms of genes differentially expressed in the mesodermal or neuronal lineages at the two different time windows.

We analysed the nuclear transcriptome by high-throughput RNA sequencing (RNA-seq) using INTACT-sorted mesodermal or neuronal nuclei obtained from stage 10 to 13 (4-9h after egg lay (AEL)) and stage 14 to 17 embryos (10-18h AEL). Robust and reproducible data were obtained for all samples in biological duplicates. In total, transcripts (containing over 10 RPKM) corresponding to 4588 coding genes were identified for the early mesodermal, 4133 coding genes for late mesodermal, 5336 coding genes for the early neuronal and 4833 coding genes for the late neuronal nuclei populations, which included genes typical for the lineage type and developmental stage (Supplementary Tab. 1). Pearson correlation coefficient analysis revealed that the transcriptome of the mesodermal lineage was clearly distinct from the neuronal one at both time points (r=0.72 for stages^10–13^; r=0.53 for stages^14–17^) (Fig. 1F, Supplementary Fig. 1I). This was also reflected in a high number of genes differentially expressed in the mesodermal as well as the neuronal lineages when comparing identical stages (Fig. 1G, H). Importantly, GO term classification revealed a significant enrichment of processes typical for the respective lineage (mesoderm both stages: p-value < 2.2e-16, neuronal stages^10–13^: p-value < 1.26e-6, neuronal stages^14–17^: p-value < 5.3e-5) (Figs 1G, 1H). In addition to elucidating differences in tissue profiles, Pearson correlation coefficient analysis also showed that global gene expression in the mesodermal lineage changed substantially over the selected time points (r=0.78 for mesoderm stages^10–13^ + stages^14–17^) (Fig. 1F, 1H, Supplementary Fig. 1I). Tissue-and stage-dependent differences and similarities were very well reflected in the distances calculated by principal component analysis (PCA) (Fig. 1E). PCA analysis also showed that in contrast to the mesoderm the neuronal transcriptomes were very similar at both developmental time frames (Supplementary Fig. 1I), which we assumed to be a consequence of the earlier onset of nervous system differentiation^27,28^.

We next profiled genome-wide Ubx binding in the same lineages and identical time windows by chromatin immunoprecipitation coupled to massively parallel sequencing (ChIP-seq) using 1×10^6^ INTACT-sorted mesodermal and neuronal nuclei and an Ubx specific antibody generated and verified in the lab (see Materials and Methods). The data was benchmarked by the identification of Ubx binding to known target loci. One example is the well-characterized interaction of Ubx with the *decapentaplegic* (*dpp*) enhancer, which is required for *dpp* activation in parasegment 7 of the embryonic visceral mesoderm^29,30^. We found Ubx to interact with the *dpp* visceral enhancer using chromatin from INTACT sorted mesodermal but not neuronal nuclei (Fig. 2A), showing that our data reflected Ubx interactions *in vivo*. However, this analysis also uncovered that Ubx bound a large fraction of genes in both lineages (Supplementary Fig. 2D, E), although this TF is known to have different functions in the developing mesoderm and nervous system^31–34^. To resolve this discrepancy, we overlapped Ubx binding events and transcriptome profiles. We found that only 19 to 27% of the Ubx chromatin interactions occurred in the vicinity of genes actively transcribed either in the mesodermal or neuronal lineages at the different stages (Fig. 2B, Supplementary Fig. 2A), with more than 80% of these interactions occurring at intron, intergenic and distal enhancer regions (Fig. 2E). This included *Mef2*, *teashirt* (*tsh*) and *β-Tubulin at 60D* (β*Tub60D*) in early and *thin* (*tn*), *α-actinin* (*Actn*) and *Tropomyosin 1* (*Tm1*) in late mesodermal nuclei, while in early neuronal nuclei *deadpan* (*dpn*), *huckebein* (*hkb*) and *Neurotrophin 1* (*NT1*) in late neuronal nuclei *Neuroglian* (*Nrg*), *target of PoxN* (*tap*) and *castor* (*cas*) was among the Ubx bound active genes. In contrast, the majority of the Ubx chromatin interactions (72 to 80%) were close to inactive genes in the two lineages (Fig. 2B, Supplementary Fig. 2A) While it is well documented that Hox TFs function as activators and repressors depending on the context^35,36^, the high number of Ubx chromatin interactions at non-transcribed genes was unexpected. By determining the gene functions associated with Ubx interactions, we found a substantial overrepresentation of GO terms characteristic for the respective lineage among the Ubx targeted and expressed genes (mesoderm: p-value 2.2e-16, neuronal stages^10–13^: p-value 3.3e-6, neuronal stages^14–17^: p-value 0.002) (Fig. 2C, D), while Ubx interactions at inactive genes occurred frequently at genes controlling processes active in other lineages (Supplementary Fig. 2A-2C). For example, ectodermal and neuronal but not mesodermal GO terms were highly enriched among the inactive genes bound by Ubx in the early mesodermal lineage (neuronal^10–13^: p-value 0.0014, ectoderm^10–13^: p-value 0.05).

**Fig. 2:**
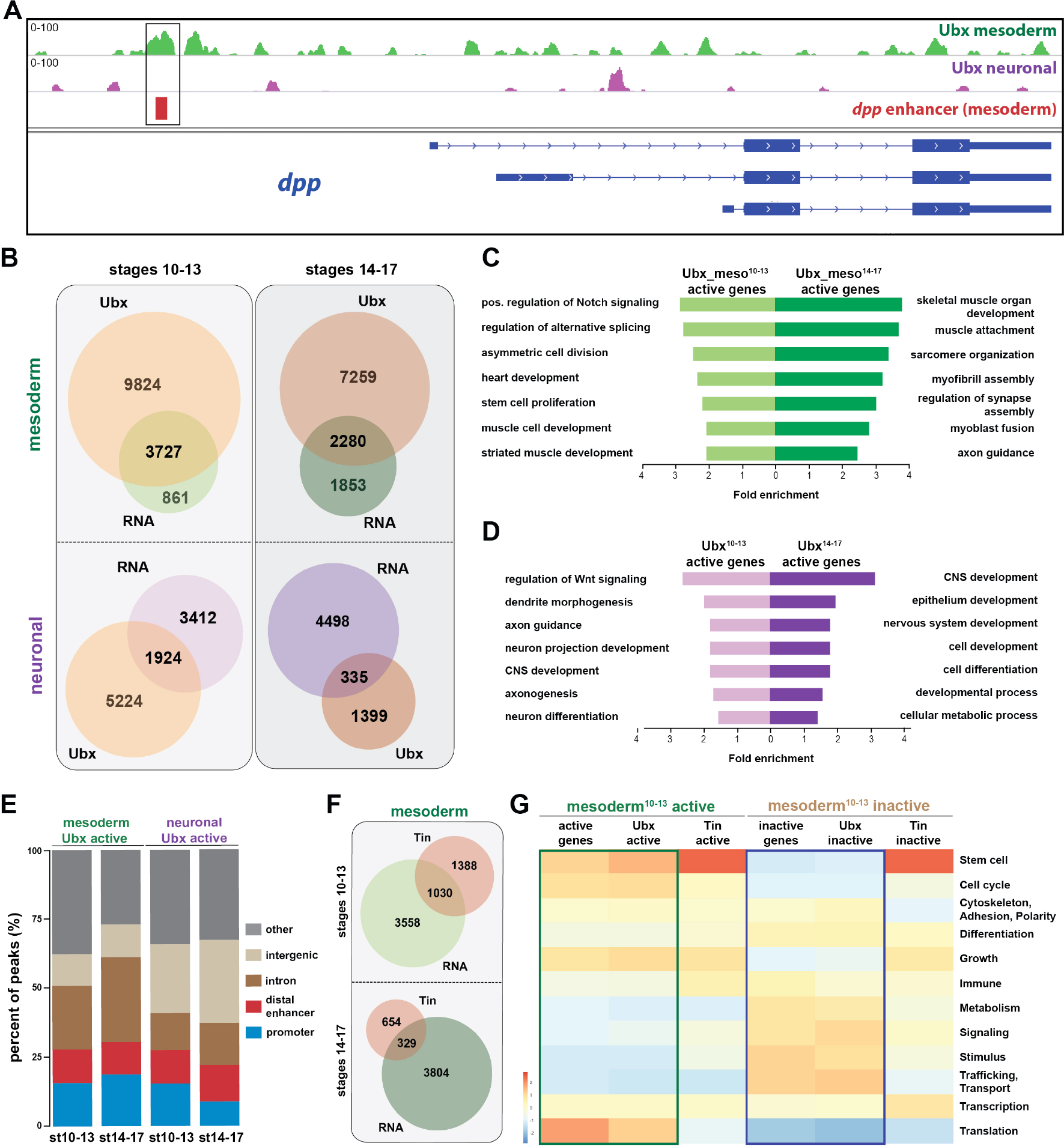
Ubx comprehensively controls tissue-specific transcriptional programs. **(A)** ChIP-seq binding profiles of Ubx at the *dpp* genomic locus in mesodermal (green) and neuronal (purple) nuclei. The different isoforms of the *dpp* gene are shown in blue, the known *dpp* visceral enhancer in red. The box highlights Ubx binding to the *dpp* visceral enhancer in mesodermal but not in neuronal cells. **(B)** Venn diagrams representing the overlaps between the mesodermal (green) and neuronal (purple) transcriptomes and Ubx bound genes (orange) at two different stages. **(C, D)** Fold enrichment of gene ontology terms of genes expressed and targeted by Ubx in the mesoderm (C) or the neuronal (D) tissues, respectively. **(E)** Comparison of the localisation of peaks called within unique genomic regions of Ubx. Locations are classified as promoters (−1000-+10 bp from TSS, 5’ UTR), distal enhancers (−2000 to −1000 from TSS, 3’ UTR, downstream), intron (intronic regions), intergenic (distal intergenic) and other regions (including exons). **(F)** Venn diagrams representing the overlaps between the mesodermal (green) transcriptomes and Tin bound genes (red) at two different stages. **(G)** Heat-map displaying presence of genes belonging to higher-order categories in the different gene classes. The colour range corresponds to the centred and scaled (per column) fraction of genes annotated to the category that also appear in the sample: red colour represents high values, blue colours low fractions of genes in the category, which are also present in the sample. Rows and columns are hierarchically clustered using Euclidean distance with complete linkage. The blue and green boxes highlight the distributions of functional terms among the expressed and Ubx targeted as well as the inactive and Ubx targeted genes.

In order to comprehensively analyse the binding behaviour of Ubx to active and inactive genes, we used the WEADE tool, which identifies and visualizes overrepresentation of functionally related biological GO terms assembled to higher-order GO term sets and allows the representation and comparative analysis of multiple gene sets ^37^. This analysis uncovered a high correlation between the lineage-specific transcriptional profiles and the genome-wide Ubx interactions (Fig. 2G). For example, many genes expressed in the early mesoderm controlled stem cell, cell cycle and translation related processes, and Ubx interactions were found enriched in the vicinity of these genes, while Ubx hardly interacted with genes encoding gene functions that were not represented among the active genes, like metabolism, signalling, stimulus and transport/trafficking related functions (Fig. 2G). A similar correlation was found among the inactive genes, as Ubx interactions were again highly enriched at gene classes found to be overrepresented among the non-expressed genes, while Ubx did not interact with underrepresented gene classes (Fig. 2G). This result suggested that Ubx binding controlled global gene expression in the mesoderm, the activation as well as repression, in a comprehensive manner. In order to test whether this binding behaviour was a general characteristic of TFs, we analysed GO term enrichment of genes bound by Tinman (Tin), a NK homeodomain TF expressed exclusively in the mesoderm^38^. To this end, we used genome-wide binding data of Tin profiled at similar embryonic stages^39^, and analysed higher-order GO term representation among the genes bound by Tin that were either transcribed or silent in the mesoderm. In contrast to Ubx, Tin interactions were mostly independent of gene classes represented among the active and inactive genes (Fig. 2G). Indeed, Tin preferentially interacted with genes controlling stem cell processes irrespective of whether these genes were actively transcribed or silent (Fig. 2G). We assumed the differential binding behaviour of Ubx and Tin to reflect the more restricted function of Tin in the mesoderm, as this TF controls the determination and specification of the cardiac, visceral and dorsal mesoderm^40,41^, while Ubx seemed to generally control development of the mesoderm (Fig. 2G).

In sum, these results illustrated that the broadly expressed Hox TF Ubx played a prominent role in orchestrating the transcriptional program in the mesodermal (and neuronal) lineages. In addition, our analysis revealed that a substantial fraction of Ubx interactions were found in the vicinity of inactive genes, which encoded many mesoderm-unrelated functions. Thus, we hypothesized that one important function of Ubx in tissue development could be the repression of alternative transcriptional programs, which instruct the development of other lineages.

### The generic Hox transcription factor Ubx functions as a major regulator of lineage programs

Our analysis of Ubx function in the mesodermal and neuronal lineages was so far based on correlating lineage-specific Ubx binding profiles with RNA-seq probed gene expression. In order to elucidate which genes are under direct Ubx control, we profiled the transcriptional output induced in mesodermal cells devoid of Ubx protein, while leaving Ubx levels in all other cell and tissue types unchanged. We focused our analysis on the mesoderm, as its control by various lineage-restricted TFs is well described^40–44^. In order to deplete Ubx in the mesoderm, we used the targeted degradation of GFP fusion proteins. To this end, we first generated an endogenously GFP tagged version of the *Ubx* gene using the Clustered Regularly Interspaced Short Palindromic Repeats-associated Cas9 (CRISPR/Cas9)-mediated genome engineering (Supplementary Fig. 3A, B)^45,46^ (see Materials and Methods for details). Intactness of the fusion protein in the homozygous viable GFP-Ubx fly line was verified by rescuing *Ubx* mutants using the *GFP-Ubx* allele (Supplementary Fig. 3H-K). Co-localization of GFP and Ubx protein expression in GFP-Ubx embryos (Fig. 3A, B, Supplementary Fig. 3C, D) confirmed a precise expression control of the fusion protein. In order to degrade the GFP-Ubx fusion protein in a lineage-specific manner, we used the deGradFP system, which harnesses the ubiquitin-proteasome pathway to achieve direct depletion of GFP-tagged proteins^20^. We first functionally verified the system by ubiquitously degrading the GFP-Ubx fusion protein using the *armadillo* (*arm*)-GAL4 driver^47^, which resulted in a strong reduction of GFP-Ubx protein levels (Supplementary Fig. 3E-G, O), and in animals resembling the *Ubx* null mutant phenotype (Supplementary Fig. 3M, n)^10^. In a next step, we applied deGradFP to specifically interfere with Ubx function in the mesoderm using the *Mef2*-GAL4 driver^48^. This combination substantially decreased GFP-Ubx protein accumulation in the mesoderm (Fig. 3C, D, G), and the expression of the direct Ubx target gene *dpp* in the visceral mesoderm (Fig. 3H-J)^30,49^. Consequently, we only observed the well-described loss of the third midgut constriction in these embryos (Fig. 3E, F), which is caused by the absence of Ubx activity in this tissue^50,51^, while all non-mesodermal tissues were unaffected (Fig. 3E-G). Having confirmed the suitability of the approach, we INTACT-sorted mesodermal nuclei, in which the GFP-Ubx protein had been depleted and profiled their transcriptome using RNA-seq. Due to the activity of the *Mef2*-GAL4 driver starting at embryonic stage 7^48^, which resulted in significant reduction of Ubx protein levels in the mesoderm only at stage 12 (Fig. 3C, D), we restricted our analysis to the late developmental time window (embryonic stages 14 to 17). Comparison of the Ubx depleted and control mesodermal transcriptomes revealed a differential expression of 2845 genes, with 1393 genes showing increased mRNA accumulation, including *Fasciclin 2* (*Fas2*), *ventral veins lacking* (*vvl*), *Neurotrophin 1* (*NT1*) and *myospheroid* (*mys*), while 1452 genes exhibited reduced expression, including *βTub60D*, *Ankyrin 2* (*Ank2*) and *Notchless* (*Nle*) (Fig. 3K). PCA analysis confirmed that the mesodermal transcriptomes in the absence (mesoderm^14–17^ Ubx^Degrad^) or presence (mesoderm^14–17^) of Ubx were substantially different (Fig. 3N). Strikingly, 85% (1227/1452) of the genes with reduced and 90% (1299/1393) of the genes with increased expression were bound by Ubx in mesodermal nuclei in wild-type embryos, implying that most of the expression changes were a direct consequence of altered Ubx chromatin interactions. To have a global view of the biological processes directly controlled by Ubx, we compared overrepresentation of higher-order biological GO terms between the Ubx degradation and control transcriptomes sets using the WEADE tool. We observed that processes were not randomly changed in the absence of Ubx but were mostly in agreement with Ubx chromatin binding (Fig. 3M). For example, translation and cell cycle processes, which were enriched among the expressed as well as Ubx bound active genes in the control transcriptome, were now represented among the transcripts with reduced accumulation in the absence of Ubx (Fig. 3M), supporting that these processes were directly activated by Ubx. On the other hand, stimulus and signalling related processes, which were only to a minor extent represented among the genes expressed in the mesoderm, were found enriched among the genes with enhanced activity in the absence of Ubx (Fig. 3M). Consistent with a repressive function of Ubx, these processes were overrepresented among the inactive genes bound by Ubx in control mesodermal nuclei (Fig. 3M).

**Fig. 3:**
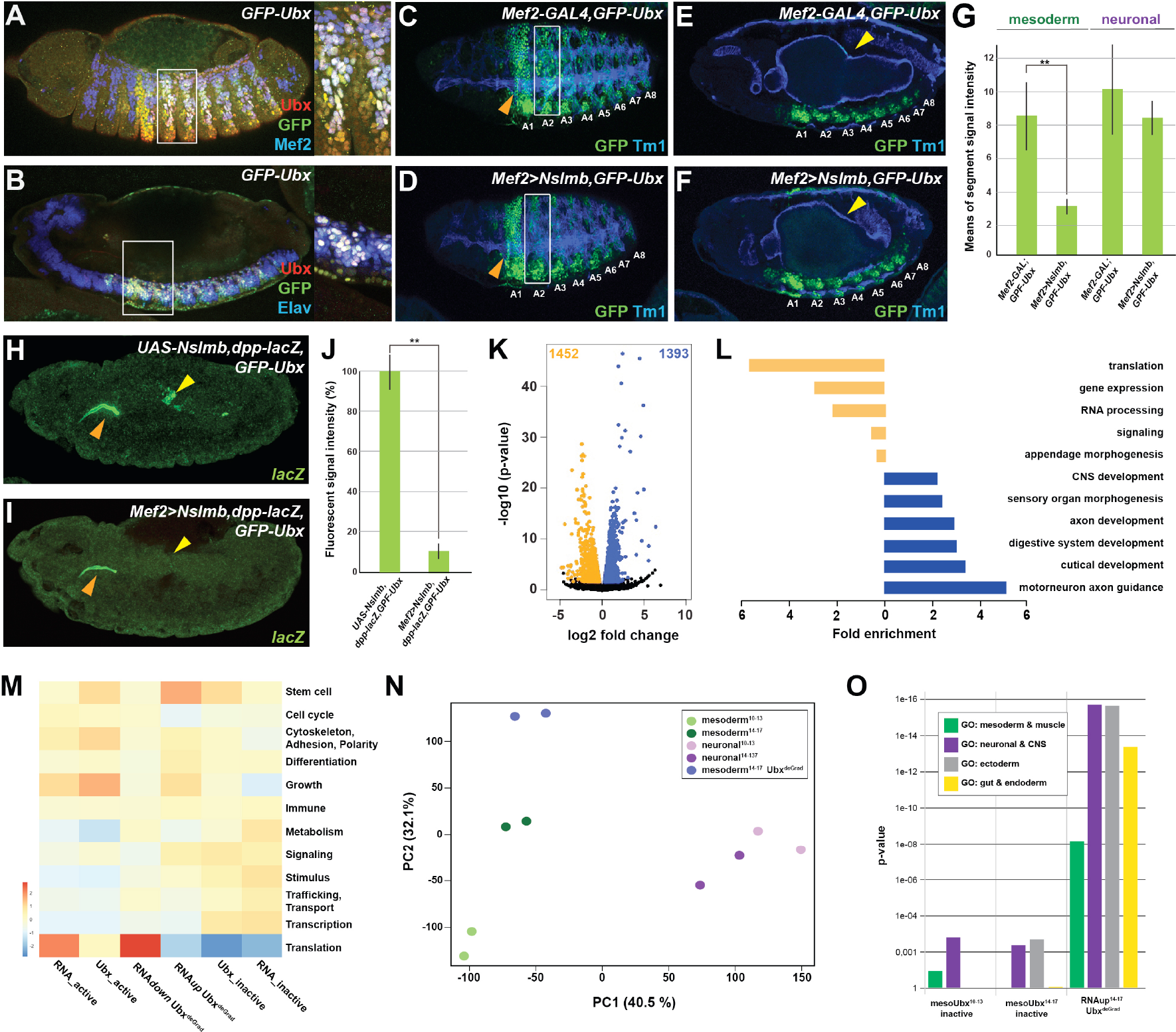
Ubx directly represses alternative fate genes in a lineage-specific manner. **(A, B)** Lateral view of stage 14 *GFP-Ubx Drosophila* embryos stained for the muscle differentiation marker Mef2 (blue) (A), the pan-neuronal marker Elav (blue) (B), Ubx (red) and GFP (green). Boxes indicate the location of the close-ups on the right panel. **(C-F)** Lateral view of stage 15 *Mef2-GAL4,GFP-Ubx* and *Mef2>Nslmb,GFP-Ubx Drosophila* embryos stained for Tm1 (blue) to indicate the differentiated muscles and GFP (green) to highlight GFP-Ubx expression. The boxes in (C, D) mark GFP expression in muscle cells of the 2^nd^ abdominal segment (A2), which is lost when Ubx is degraded, while the ectodermal expression in the 1^st^ abdominal segment (A1) is unaffected (marked by orange arrowheads). In (E, F), GFP expression in the CNS and the visceral mesoderm (marked by yellow arrowheads) is shown. GFP expression in the visceral mesoderm is lost in *Mef2>Nslmb,GFP-Ubx Drosophila* embryos (F), leading to a loss of the second midgut constriction. **(G)** Quantification of GFP signal intensity in the mesoderm and the CNS of *Mef2-GAL4,GFP-Ubx* and *Mef2>Nslmb,GFP-Ubx Drosophila* embryos, showing that GFP is strongly decreased in the mesoderm (** = p<0.01). **(H, I)** *lacZ* mRNA expression in stage 14 *UAS-Nslmb,dpp-lacZ,GFP-Ubx* and *Mef2>Nslmb,dpp-lacZ,GFP-Ubx Drosophila* embryos, highlighting that mesoderm-specific depletion of Ubx leads to a loss of *dpp-lacZ* enhancer activity (indicated by yellow arrowheads), a known and direct target of Ubx control^29,30^. The orange arrowheads highlight unspecific enhancer activity in the salivary glands. **(J)** Quantification of *lacZ* signal intensity in *UAS-Nslmb,dpp-lacZ,GFP-Ubx* and *Mef2>Nslmb,dpp-lacZ,GFP-Ubx* embryos, showing that *lacZ* expression is strongly decreased (** = p<0.01). **(K)** Volcano plot displaying the differentially expressed genes between *UAS-Nslmb,GFP-Ubx* and *Mef2>Nslmb,GFP-Ubx* INTACT-sorted mesodermal nuclei. The y-axis corresponds to the mean expression value of log10 (p-value), and the x-axis displays the log2 fold change value. The orange dots represent transcripts whose expression is down-regulated (padj-value < 0.1), the blue dots represent the up-regulated expressed transcripts (padj-value < 0.1) between *UAS-Nslmb,GFP-Ubx* and *Mef2>Nslmb,GFP-Ubx* INTACT-sorted mesodermal nuclei. **(L)** Fold enrichment of gene ontology terms of down-(orange) and up-regulated (blue) genes in *Mef2>Nslmb,GFP-Ubx* vs. *UAS-Nslmb,GFP-Ubx* mesodermal nuclei. **(M)** Heat-map displaying presence of genes belonging to higher-order categories in the different gene classes. The colour range corresponds to the centred and scaled (per column) fraction of genes annotated to the category that also appear in the sample: red colour represents high values, blue colours low fractions of genes in the category, which are also present in the sample. Rows and columns are hierarchically clustered using Euclidean distance with complete linkage. **(N)** PCA applied to all RNA-seq samples identifies the separation of the Ubx^deGrad^ dataset from the two mesodermal as well as the neuronal datasets. **(O)** Multiple testing of higher-order GO-terms related to different lineages among different gene classes. The y-axis corresponds to the p-value, the x-axis displays the different categories of tested genes. Neuronal and ectodermal as well as gut/endoderm related GO-terms are significantly enriched in the tested samples.

Intriguingly, we found differentiation processes to be enriched among the genes up-regulated in the absence of Ubx (Fig. 3L, M). Analysing differentiation processes revealed that processes controlling the development of lineages other than the mesoderm, including the neuronal, ectodermal end endodermal lineages, were predominantly represented among the up-regulated genes in Ubx depleted mesodermal cells (Fig. 3L, Supplementary Fig. 3P), which was significantly higher than expected by chance (neuronal and ectoderm: p-value < 2.2e-16, endoderm & gut structures: p-value 5.04e-14). Comparing these genes to the neuronal transcriptomes showed that 43% (578/1392) of the up-regulated genes were normally expressed and functional in the neuronal lineage in the wild type situation (Supplementary Fig. 3P), while the remaining 57% (814/1392) were active in other lineages according to GO term prediction (data not shown).

Taken together, these results showed that Ubx controlled a large number of genes encoding diverse functions. The strong correlation between Ubx binding with specifically repressed or activated classes of genes suggested that Ubx indeed exerts a dominant influence over the mesodermal transcriptome at the two stages analysed. In addition, this analysis highlighted that Ubx repressed many genes controlling the establishment of non-mesodermal, alternative lineages, indicating that Ubx could have a pivotal role in restricting cellular plasticity, thereby ensuring that lineages adopt a unique identity.

### Ubx represses alternative fate genes by organizing the epigenetic landscape

One question emerging from these results was how Ubx mediated the repression of non-mesodermal fate genes. As gene expression critically depends on the epigenetic status of the gene regulatory control regions^52,53^, we characterized Ubx chromatin interactions in more detail. To this end, we mapped two key modifications of the nucleosome component Histone H3, H3K27me3, a mark associated with repressive chromatin^54,55^, as well as H3K27ac, a mark associated with active promoters and enhancers^56,57^, by ChIPseq using INTACT-sorted mesodermal nuclei. This analysis revealed that in the early mesoderm about 30% of all Ubx chromatin interactions overlapped with H3K27me3 peaks and about 20% with H3K27ac peaks, while in the late mesoderm 40% of all Ubx chromatin interactions overlapped with H3K27me3 peaks and 73% with H3K27ac peaks (Supplementary Fig. 4B). Ubx interactions with repressive chromatin were predictive of gene activity, as about 80% of these occurred at inactive genes irrespective of the developmental stage (Fig. 4B), which we found again associated with functions unrelated to the mesoderm (neuronal: p-value 0.003 – 1.89e-9, ectoderm: p-value 0.006 – 4.46e-11, endoderm & gut structures: p-value 0.012) (Fig. 4C). In contrast, less than half of the Ubx interactions with active chromatin occurred close to active genes (Fig. 4B), which were as before enriched for mesoderm related functions (Fig. 4C), while the rest were found close to inactive genes (Fig. 4B). Organizing chromatin environment by genomic location revealed that only a minor fraction (about 20%) of the Ubx bound repressive chromatin regions occurred at promoters, while the majority was found at putative distal enhancers, in introns and intergenic regions in both developmental time points (Supplementary Fig. 4D).

**Fig. 4:**
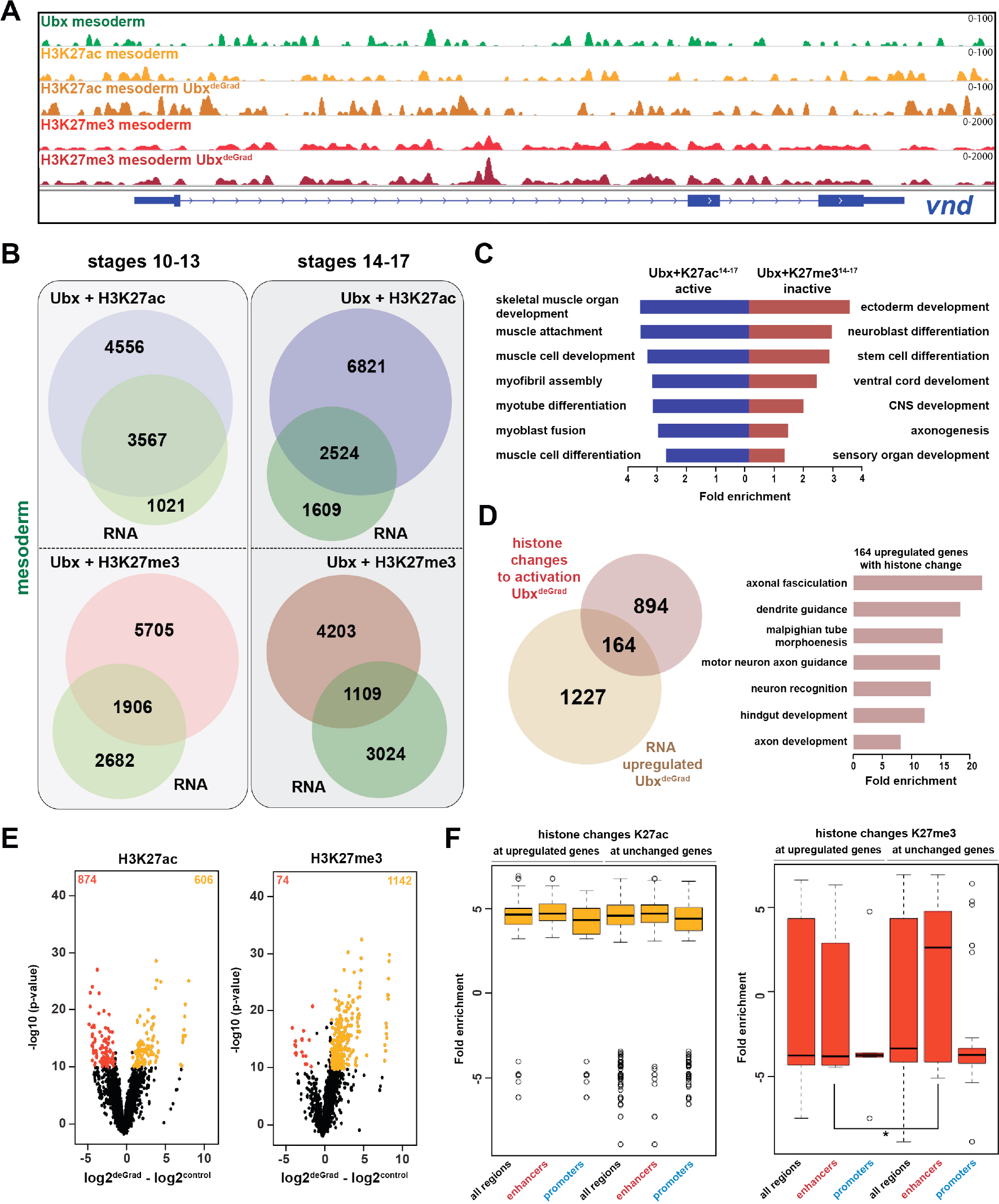
Ubx represses alternative fate genes by organizing the epigenetic landscape. **(A)** ChIP-seq binding profiles of Ubx (green), H3K27ac (light + dark orange) and H3K27me3 (light + dark red) in control (*UAS-Nslmb,GFP-Ubx*) (light orange + red) and Ubx^DeGrad^ (*Mef2>Nslmb,GFP-Ubx*) (dark orange + red) mesodermal nuclei at the *vnd* genomic locus (blue), a gene up-regulated in the absence of Ubx. **(B)** Venn diagrams representing the overlaps between genes expressed in the mesoderm (green), which are targeted by Ubx and simultaneously associated with H3K27ac (Ubx+H3K27ac, purple) or H3K27me3 (Ubx+H3K27me3, red) marks, respectively. **(C)** Fold enrichment of gene ontology terms among the genes expressed, targeted by Ubx and associated with H3K27ac marks (Ubx+K27ac^14–17^ active, blue) and the genes not expressed in the mesoderm, targeted by Ubx and associated with H3K27me3 marks (Ubx+K27me3^14–17^ inactive, red). **(D)** Left panel: Venn diagram representing the overlap between genes up-regulated in Ubx depleted mesodermal nuclei (beige) and genes associated with histone mark changes indicative for gene activation (gain/increase in H3K27 acetylation, loss/reduction in H3K27 tri-methylation) (brown). Right panel: Fold enrichment of gene ontology terms of the 164 genes, which have histone changes to activation and are up-regulated in Ubx depleted mesodermal nuclei. **(E)** Volcano plot displaying the changes of H3K27ac and H3K27me3 marks between *UAS-Nslmb,GFP-Ubx* (control) and *Mef2>Nslmb,GFP-Ubx* (degrade) INTACT-sorted mesodermal nuclei using differential binding analysis of ChIP-Seq peaks (DiffBind). The y-axis corresponds to the mean expression value of log10 (p-value), and the x-axis displays the difference of the log2 fold change value in Ubx^Degrad^ compared to the log2 of the control. The orange dots represent genomic regions containing an increase in H3K27ac or H3K27me3 (p-value < 0.05), the red dots represent genomic regions containing an decrease in H3K27ac or H3K27me3 (p-value <0.05) between *UAS-Nslmb,GFP-Ubx* and *Mef2>Nslmb,GFP-Ubx* INTACT-sorted mesodermal nuclei. **(F)** Comparison of H3K27ac or H3K27me3 fold enrichments at Ubx targeted genomic regions that experienced histone changes towards activation represented as Box plots. Quantification of the histone changes in Ubx bound upregulated (164) and unchanged genes (894) by using the results of the differential binding analysis of ChIP-Seq peaks. The y-axis displays the fold change calculated by DiffBind. The x-axis shows all selected regions (upregulated and unchanged) and a sub-division of the regions in promoter (−1000 to +10 from TSS, 5’ UTR) and enhancer (distal enhancers (−2000 to −1000 from TSS), 3’ UTR, downstream, intronic regions, distal intergenic). Each sample contains a significant increase in H3K27ac levels (yellow), H3K27me3 levels (red) are decreased, but enhancer regions close to genes not changed in their expression have significantly higher H3K27me3 levels than enhancers close to genes up-regulated after Ubx depletion (* = p<0.05).

In a next step, we asked how the epigenetic landscape changed in the absence of Ubx, in particular at genomic regions with overlapping Ubx and repressive H3K27me3 marks. In total, 1216 H3K27me3 peaks and 1480 H3K27ac peaks located in gene regulatory regions, which corresponded to 2768 genes, changed their chromatin state in the absence of Ubx, as they experienced either a reduction/loss or an increase/gain in tri-methylation/acetylation at lysine 27 of histone 3 (Fig. 4E, Supplementary Fig. 4E). Importantly, 80% of these genomic regions possessed an Ubx binding peak in control mesodermal cells, indicating that Ubx was critically required for establishing or maintaining the majority of these histone marks. Intriguingly, 1058 of these genes were primed for gene expression, as they experienced a loss/reduction of H3K27me3 and/or gain/increase of H3K27ac in their regulatory regions (Fig. 4D), and 164 of them were indeed up-regulated in the mesodermal lineage in the absence of Ubx (Fig. 4D, Supplementary Fig. 4E). Consistent with previous results, these 164 genes were highly enriched for processes critically controlling the development of lineages other than the mesoderm (neuronal: p-value < 2.2e-16, ecoderm: p-value 1.36e-12, endoderm & gut structures: p-value < 2.2e-16) (Fig. 4D).

One result that puzzled us was that the majority of genes primed for activation (894 of 1058) remained silent on the RNA level when Ubx was lineage-specifically degraded (Fig. 4D). In order to resolve this discrepancy, we analysed H3K27me3 and H3K27ac changes at introns, intergenic and distal enhancer regions, which we collectively labelled enhancers (Fig. 4F), as well as at promoters. We found that H3K27ac marks were equally increased at promoters and enhancers of genes unchanged or up-regulated in the absence of Ubx (Fig. 4F). However, while H3K27me3 marks were significantly decreased at enhancers as well as promoters of up-regulated genes, unchanged genes showed a significant increase in H3K27me3 at their enhancers (Fig. 4F). As a high proportion of Ubx and H3K27me3 peaks co-occurred at enhancers (Supplementary Fig. 4D), this result revealed that Ubx played a major role in repressing the expression of alternative fate genes by controlling the deposition/maintenance of H3K27me3 marks at enhancers.

In sum, these results showed that Ubx mediated the repression of genes encoding non-mesodermal functions by controlling the epigenetic status, in particular H3K27me3, at Ubx bound chromatin sites located in enhancers.

### Ubx interacts with Pho at H3K27me3 marked inactive genes encoding alternative fate genes

To identify factors that together with Ubx could mediate the repression of alternative fate genes thereby restricting lineage identity, we performed a DNA motif search using all Ubx peaks either overlapping with H3K27me3 repressive or H3K27ac active marks. We found in both cases motifs for Ubx, Extradenticle (Exd), a TALE class homeobox TF functioning as Hox cofactor in invertebrates and vertebrates^58,59^, and Trithorax-like (Trl), a GAGA factor activating and repressing gene expression by chromatin modification^52^, among the highest ranking motifs. In contrast, the DNA binding motif for the zinc finger protein Pleiohomeotic (Pho) was specifically enriched only among the co-occurring Ubx and H3K27me3 binding events (Fig. 5A). Interestingly, it has been shown just recently that in the absence of Pho, which recruits PcGs to PREs^60–62^, H3K27me3 marks were reduced in Polycomb regions and redistributed to heterochromatin^63^. In combination with our finding that H3K27me3 levels were reduced specifically at up-regulated genes (Fig. 4F), we hypothesized that Pho could function together with Ubx in the lineage-specific repression of alternative fate genes. Consistent with this idea Pho was found expressed in mesodermal cells also expressing Ubx (Fig. 5B – C”). Furthermore, we confirmed an interaction of Ubx and Pho proteins in a complex *in vitro* and *in vivo* by performing co-immunoprecipitation (Co-IP) experiments *in cellulo* using *Drosophila* S2R+ cells transfected with tagged versions of Ubx and Pho as well as in *vivo* using GFP-Ubx embryos (Fig. 5F, G). This result suggested that Ubx and Pho could interact with the same chromatin regions to control gene expression, thus we analysed high-resolution Pho maps retrieved from embryonic (stage 9 to 12) mesodermal cells^64^. We found 4814 chromatin regions (which represent 62% of the Ubx and 12% of the Pho binding events) to be co-bound by Ubx and Pho and marked by H3K27me3 in stage 14-17 mesodermal cells (Fig. 5D, 6a, Supplementary Fig. 5A), with 40% of these events occurring at promoters and 49% at enhancers (Supplementary Fig. 5C). Consistent with the reported role of Pho in transcriptional repression^60–62^, 76% (3578/4715) of the genes bound by Ubx and Pho as well as marked by H3K27me3 were not expressed in the mesoderm (Fig. 5D), and GO term analysis revealed that the majority of the genes encoded non-mesodermal and stem cell-related functions (p-value 2.87e-11) (Fig. 5E). This was different for the remaining 1137 genes also bound by Ubx and Pho and marked by H3K27me3 but expressed in mesodermal cells, as they encoded gene functions controlling processes typical for the tissue type and developmental stage (Fig. 5E). By analysing the distribution of H3K27me3 and H3K27ac at regulatory regions, we discovered that those chromatin regions associated with inactive genes had a higher coverage of H3K27me3 and lower coverage of H3K27ac at shared Ubx/Pho binding regions, while it was the opposite for active genes (Fig. 5I). Furthermore, the canonical Pho binding motif was found overrepresented only among the Ubx/Pho/H3K27me3 chromatin regions associated with genes inactive in the mesoderm (Fig. 5H), while other DNA binding motifs, including the ones for Ubx, Exd and Trl, were overrepresented in Ubx/Pho/H3K27me3 chromatin regions associated with inactive as well as active genes.

**Fig. 5:**
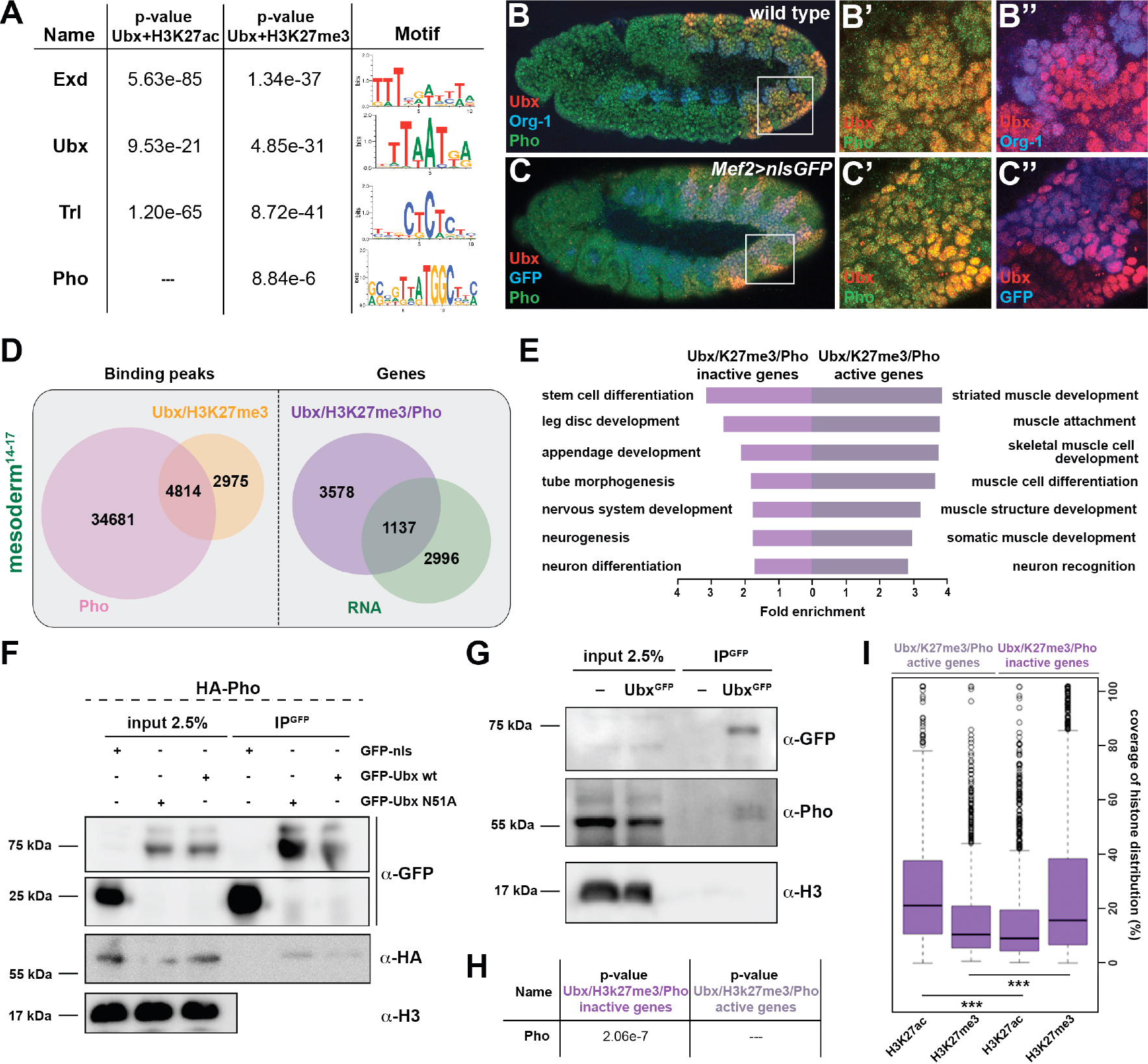
Ubx interacts with Pho at H3K27me3 marked inactive genes encoding alternative fate genes. **(A)** The top four hits from MEME-ChIP analysis using all Ubx targeted, H3K27ac marked genomic regions in the vicinity of expressed genes (Ubx+H3K27ac) or Ubx targeted, H3K27me3 marked genomic regions in the vicinity of inactive genes (Ubx+H3K27me3) as input. While the Ubx, Exd and Trl motifs were found over-represented in both datasets, the Pho motif was found over-represented only in the Ubx+H3K27me3 dataset. **(B-B’’)** Lateral view of stage 11 wild-type *Drosophila* embryos stained for the mesodermal marker Org-1 (blue), Ubx (red) and Pho (green). (B’) and (B’’) are dual colour images of (B), highlighting that Ubx and Pho are co-expressed in Org-1 labelled mesodermal cells. **(C-C’’)** Lateral view of stage 11 *Mef2>nlsGFP Drosophila* embryos stained for GFP (blue), Ubx (red) and Pho (green). (C’) and (C’’) are dual colour images of (C). **(D)** Left: Venn diagram representing the overlap between Ubx targeted and H3K27me3 marked chromatin regions (orange) and Pho binding peaks (pink) in stage 14-17 mesodermal nuclei. Right: Venn diagram representing the overlap between genes targeted by Ubx + Pho and marked by H3K27me3 (purple) and the genes expressed in the mesoderm (green). **(E)** Fold enrichment of gene ontology terms of the genes targeted by Ubx and Pho, which are additionally marked by H3K27me3 and either not expressed (3578) or expressed (1137) in mesodermal cells. **(F)** Co-immunoprecipitation of exogenous GFP fusion proteins using GFP-trap beads after transfection of S2R+ *Drosophila* cells with HA-Pho coupled with GFP-nls (negative control), GFP-Ubx WT or N51A (mutant of the homeodomain, Asparagine 51 replaced by Alanine residue). Western blots were probed with the indicated antibodies. Pho is detected in the immunoprecipitated fraction of GFP-Ubx WT and N51A, while it is absent in the GFP negative control. **(G)** Co-immunoprecipitation with the GFP-trap system of endogenous proteins performed on nuclear extract from embryos expressing (or not) the endogenous GFP-Ubx fusion protein. Pho co-immunoprecipitates with GFP-Ubx, while it is absent from w1118 fly line. **(H)** MEME-ChIP analysis using Ubx and Pho targeted chromatin regions marked by H3K27me3 located in the vicinity of active or inactive genes identifies the classical Pho motif over-represented only in the vicinity of inactive genes. **(I)** Quantification of H3K27ac and H3K27me3 distributions at Ubx and Pho targeted chromatin regions marked by H3K27me3 located either in the vicinity of active or inactive genes are represented as Box plot. The y-axis displays the coverage of the histone mark distribution in percent, the x-axis indicates the histone mark that was analysed. Active genes display a higher coverage of H3K27ac, while inactive genes have a higher coverage of H3K27me3 (*** = p<0.001).

**Fig. 6:**
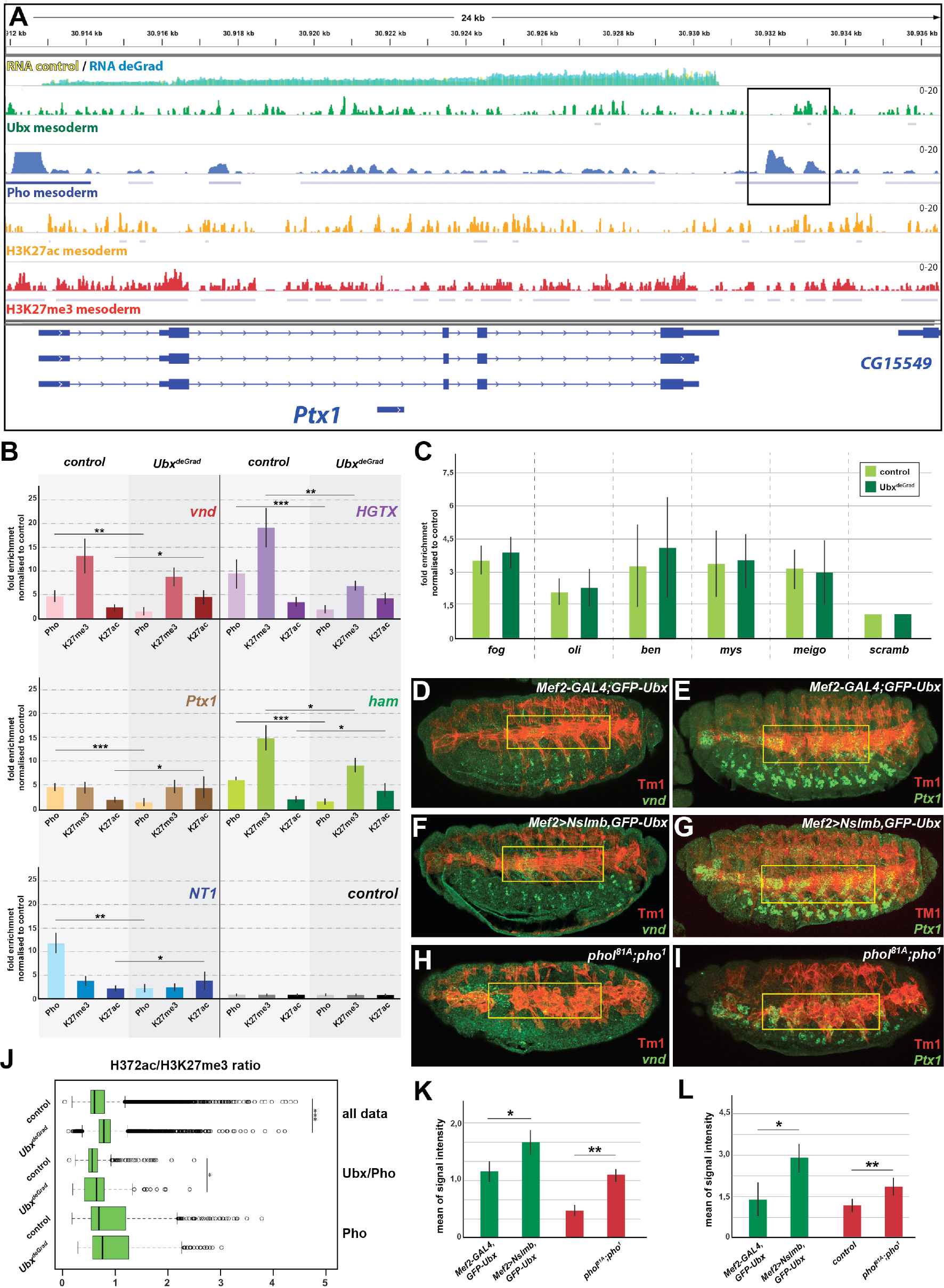
Ubx is required for stabilizing Pho binding to H3K27me3 chromatin regions. **(A)** ChIP-seq binding profiles of Ubx (green), Pho (Pho), H3K27ac (orange) and H3K27me3 (red) in mesodermal nuclei as well as RNA-seq profiles in *Mef2-GAL4,GFP-Ubx* control (yellow) versus and *Mef2>Nslmb,GFP-Ubx* (blue) mesodermal nuclei at the *Ptx1* genomic locus. The *Ptx1* gene are shown in blue. The box highlights the overlapping binding peaks of Ubx and Pho in the vicinity of the *Ptx1* coding region. **(B)** ChIP-qPCR experiments for Pho, H3K27ac and H3K27me3 using chromatin regions close to the *vnd*, *HGTX*, *Ptx1*, *ham* and *NT1* genes bound by Pho and Ubx using chromatin isolated from *UAS-Nslmb,GFP-Ubx* (light green) and *Mef2>Nslmb,GFP-Ubx* (dark green) mesodermal nuclei. As control locus sites, a exon region in the *scramb* genes was used, which is not bound by Pho nor by Ubx. All loci show a significant reduction of Pho binding, H3K27me3 levels were reduced at the HGTX associated Ubx/Pho chromatin regions. In the *vnd*, *Ptx1*, *ham* and NT1 associated Ubx/Pho loci H3K27ac levels are significantly enriched (* = p<0.05, ** = p<0.01, *** = p<0.001). **(C)** ChIP-qPCR experiments of Pho detecting chromatin regions close to the *fog*, *oli*, *ben*, *mys* and *meigo* genes bound by Pho but not Ubx and chromatin isolated from *UAS-Nslmb,GFP-Ubx* (light green) and *Mef2>Nslmb,GFP-Ubx* (dark green) mesodermal nuclei. As control locus, a exon region in the *scramb* genes was used, which is not bound by Pho or Ubx. **(D-I)** Lateral view of stage 16 *Mef2-GAL4,GFP-Ubx* (D, E), *Mef2>Nslmb,GFP-Ubx* (F, G) and *phol^81A^;pho^1^* mutant (H, I) embryos stained for the muscle marker Tm1 (D-I) and for *vnd* (D, F, H) and *Ptx1* (E, G, I) transcripts. The yellow boxes highlight the lateral muscles. **(J)** Analysis of acetylation (H3K27ac) to methylation (H3K27me3) ratio of the *UAS-Nslmb,GFP-Ubx* (control) and *Mef2>Nslmb,GFP-Ubx* (deGrade) using the entire dataset (full data), all reads bound by Ubx and Pho that contain histone modifications towards activation (Ubx/Pho) and reads only bound by Pho that are associated with methylation (Pho). The y-axis displays the fold change ratio of H3K27ac and H3K27me3. The x-axis shows the selected regions. The entire dataset as well as the reads containing Ubx and Pho co-bound regions indicate a higher ratio in the absence of Ubx compared to the control. The regions that are only bound by Pho show no difference in the absence of Ubx compared to the control (*** = p<0.001, * = p<0.05). **(K, L)** Quantification of *vnd* (K) and *Ptx1* (L) signal intensities in the lateral muscles (as indicated by the yellow boxes) in *Mef2-GAL4,GFP-Ubx*, *Mef2>Nslmb,GFP-Ubx* and *phol*^*81A*^;*pho*^1^ mutant embryos, showing that the expression of *vnd* and *Ptx1* is significantly increased in comparison to control embryos (* = p<0.05).

These results demonstrated that Ubx interacted with Pho and that their interaction on H3K27me3 marked chromatin regions occurred preferentially at inactive genes in mesodermal cells.

### Ubx is required for stabilizing Pho binding to H3K27me3 chromatin regions

Our results indicated that Ubx could mediate the repression of alternative fate genes by recruiting or stabilizing Pho binding at Ubx targeted genomic regions in the mesodermal lineage, thereby allowing the two proteins to act in a combinatorial fashion. In support of this hypothesis, we found 55% (769/1391) of the genes up-regulated in the absence of Ubx, which were highly enriched for non-mesodermal and stem cell related functions, to be located in the vicinity of Ubx/Pho binding regions (Supplementary Fig. 6B). In contrast, only a minor fraction (15%) was in the vicinity of chromatin sites targeted by Pho only, and we did not detect any significant enrichment of GO terms among these genes (Supplementary Fig. 6B). To provide additional evidence for a combined action of Ubx and Pho, we analysed the binding of Pho to Ubx/Pho/H3K27me3 binding regions in the absence of Ubx by performing ChIP experiments on INTACT sorted control and Ubx depleted mesodermal nuclei. Furthermore, we also quantified the levels of H3K27me3 and H3K27ac marks at these loci. We selected five Ubx/Pho/H3K27me3 genomic regions that changed their histone marks towards activation (less H3K27me3 and/or more H3K27ac) when Ubx was depleted in the mesodermal lineage, which resulted in the activation of the associated genes encoding neuronal functions. This included *ventral nervous defective* (*vnd*), a NK2 class TF encoding gene critical for patterning of the neuroectoderm as well as the formation and specification of ventral neuroblasts^65^, *HGTX*, a homeodomain TF encoding gene promoting the specification and differentiation of motor neurons innervating the ventral body wall muscles^66^, *Neurotrophin 1* (*NT1*), a cytokine encoding a gene that regulates motor neuron survival and axon guidance^67^, *hamlet* (*ham*), a gene encoding a PRDM class TF that regulates neuron fate selection in the peripheral nervous system^68^, and *Ptx1*, again a homeodomain TF encoding gene which is expressed at high levels in the early embryonic central nervous system and the midgut, and later also in ventral muscles ^69^. In addition, we also chose five loci bound by Pho but not by Ubx, which were associated with genes normally not expressed in the mesoderm and which remained silent in the absence of Ubx, including *folded gastrulation* (*fog*), *Olig family* (*Oli*), *bendless* (*ben*), *myospheroid* (*mys*) and *medial glomeruli* (*meigo*). Strikingly, all five Ubx/Pho/H3K27me3 loci, which were all co-bound by Ubx and Pho in the presence of Ubx (Supplementary Fig. 5A), experienced a dramatic loss of Pho binding when Ubx was degraded (Fig. 6B). In contrast, Pho binding to Pho-only control loci remained unaffected (Fig. 6C). As Pho expression levels were unaltered in the absence of Ubx (Supplementary Fig. 5B), we concluded that Ubx is required to stabilize Pho binding to chromatin of H3K27me3 marked loci. These results indicated that the expression of alternative fate genes should be similarly de-repressed in the mesoderm in the absence of either Ubx or Pho. In line with this hypothesis, we found the expression of *vnd* and *Ptx1*, which were normally not or only weakly expressed in mesodermal cells of stage 16 control embryos (Fig. 6D, E) to be significantly increased in the mesoderm of *Mef2>Nslmb*,*GFP-Ubx* embryos (Fig. 6F, G, K, L, Supplementary Fig. 6C-E) as well as in *phol*^*81A*^/*pho*^*1*^ double mutant embryos (Fig. 6H, I, K, L).

It has been recently reported that in the absence of Pho H3K27me3 enrichment was decreased^63,70^, and consistently, we found that H3K27me3 levels at the *HGTX* and *ham* associated Ubx/Pho/H3K27me3 loci were significantly decreased in Ubx depleted mesodermal nuclei (Fig. 6B). Interestingly, while H3K27ac levels remained unaffected at the *HGTX* associated Ubx binding locus, they were increased at the *ham* associated Ubx/Pho/H3K27me3 region (Fig. 6B). In contrast, H3K27me3 levels at Ubx/Pho/H3K27me3 loci associated with the *vnd*, *NT1* and *Ptx1* genes were not considerably decreased, however, these loci had significantly higher H3K27ac levels in the absence of Ubx (Fig. 6B). These results revealed that in the absence of Ubx Pho’s ability to interact with the five Ubx/Pho/H3K27me3 loci was strongly reduced and concomitantly H3K27me3 as well as H3K27ac levels were changed. Thus, we asked whether H3K27me3 and H3K27ac levels were generally altered at Ubx/Pho binding sites when Ubx was depleted in the mesodermal lineage. To this end, we calculated H3K27ac/H3K27me3 ratios at all Ubx/Pho/H3K27me3 loci as well as at loci only bound by Pho. Intriguingly, we found that H3K27ac/H3K27me3 ratios were significantly higher at Ubx/Pho/H3K27me3 loci in Ubx depleted mesodermal nuclei when compared to control nuclei, while they were not significantly changed at Pho-only sites (Fig. 6J).

In sum, these results demonstrated that Ubx stabilized Pho binding to chromatin regions at alternative fate genes in the mesodermal lineage, which controlled the proper levels of H3K27ac to H3K27me3 at these sites, thereby ensuring repression of these genes.

## DISCUSSION

Cell and tissue types get different during development and their identity needs to be maintained also in adulthood to guarantee the survival of organisms. Here, we provide evidence that multi-lineage TFs of the Hox class stabilize the different lineage choices by restricting cellular plasticity in a lineage specific manner. To this end, we studied the broadly expressed Hox TF Ubx in the mesodermal and neuronal lineages during *Drosophila* development using a comparative genomic approach and an experimental system to deplete Ubx protein exclusively in the embryonic mesodermal lineage. This approach allowed us to dissect the cell-autonomous function of Ubx in a single lineage that was located in an otherwise normal cellular environment at the transcriptome and chromatin level. Using this experimental set-up, we found that Ubx comprehensively orchestrates the transcriptional programs of the mesodermal as well as of the neuronal lineage, as it bound and regulated a substantially fraction of genes specifically expressed in these tissue lineages. Strikingly, this analysis revealed that the majority of Ubx chromatin interactions were located in the vicinity of inactive genes, and lineage-specific interference with Ubx in the mesoderm demonstrated that about 20% of these interactions were important for repressing the close-by genes. Intriguingly, these genes were highly enriched for alternative cell fates, demonstrating that Ubx had indeed a pivotal role in restricting developmental plasticity in a context-dependent manner.

One important question arising from this result was why not more of the inactive genes bound by Ubx were de-repressed in the absence of Ubx. There are several explanations for this finding. First, Hox TFs cross-regulate each other’s expression, a phenomenon described as posterior suppression^71,72^. Consistently, the *Hox* gene *Antennapedia* (*Antp*), which is normally expressed anterior to Ubx, is ectopically activated when Ubx function is absent (Supplementary Fig. 7), allowing Antp now to partially take over the function of Ubx in this lineage. Second, gene regulation is tightly linked to the chromatin status at promoter and enhancer regions. It had been shown in mammalian cells that the turn-over rates of the histone variant H3.3 at regulatory regions were correlated with specific histone modifications, high turn-over when associated with high levels of active histone modifications, like H3K27ac, while much slower turn-over when associated with higher levels of H3K27me3 marks^70^. This implies that the “clearing” of repressive histone modifications takes much longer in comparison to active ones. Third, H3K27me3 marks serve as epigenetic memory to permanently silence genes in the course of development, and a recent study demonstrated that a resetting of the epigenetic status requires cell division to dilute the H3K27me3 mediated silencing effect ^73^. In line with these studies, we found that the reduction of H3K27me3 levels was low in comparison to the increase of H3K27ac levels in Ubx depleted mesodermal nuclei. Thus, a de-repression of genes might require either more time or cell divisions or both. However, after its specification at embryonic stage 12 the mesoderm does not divide anymore, which might prevent an efficient clearing of H3K27me3 marks at Ubx targeted chromatin sites. Fourth, de-repression of genes is not only a consequence of abolishing repression but also of gaining activation, which not only requires a change of the histone environment at control regions but also the expression and action of the proper sets of TFs. Comparing the transcriptomes of the mesodermal and neuronal lineages revealed that only about 5% of the TFs expressed in an alternative lineage, in this case the neuronal one, were expressed in the Ubx depleted mesodermal cells, including vnd, HGTX, ham and Ptx1, which was obviously not sufficient to induce a lineage switch, in particular as the expression of only about 10% of the mesodermal fate related TFs was reduced in their expression. As we find Antp to be ectopically expressed in the mesodermal lineage in the absence of Ubx (Supplementary Fig. 7), we assume Antp to partially take over the lineage-specific function of Ubx. Thus, it will be interesting in future to study mesoderm development in a Hox-free environment and determine the fate of the developing cell lineage.

**Fig. 7:**
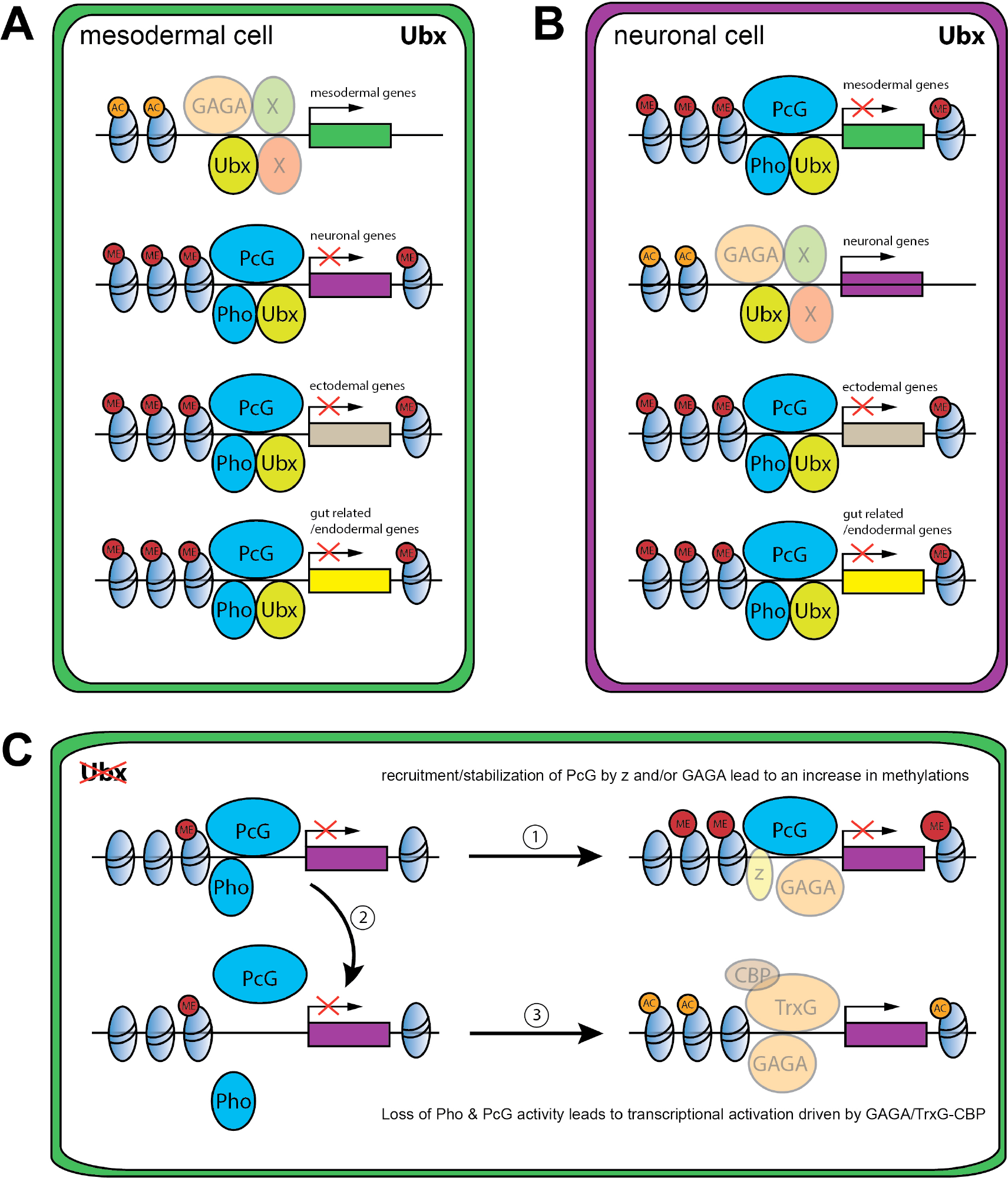
Model of Ubx and Pho combinatorial action in mesoderm development. **(A, B)** In wild-type cells of the mesodermal (A) or neuronal lineage (B) (Ubx+), Ubx (light green) is involved in the activation of mesodermal (green) or neuronal (purple) genes, possibly together with Trl (GAGA) and/or other so far unknown proteins. In the same cells, Ubx stabilizes Pho and PcG binding (dark blue) to alternative fate genes (grey: ectodermal genes, yellow: gut related genes) to ensure their lineage-specific repression. Histone tri-methylation marks (H3K27me3, red, ME) are set and maintained. Histones are illustrated in light blue and H3K27ac mark is coloured in orange (AC). **(C)** Loss of Ubx for example in the mesodermal lineage destabilises Pho binding to regulatory regions of alternative fate genes (example for neuronal (purple)), which could have different outcomes: (1) The PcG complex might be further maintained at these sites through Trl (GAGA), an interactions potentially leading to an increase of H3K27me3 levels. (2) Destabilisation of Pho and PcG at the regulatory regions of neuronal genes (purple). (3) Loss of PcG might increase TrxG binding through Trl (GAGA) in combination with CBP. This interaction could mediate the acetylation of Lys27 at histone 3 (H3K27ac, orange), resulting in the activation of alternative fate genes (example for neuronal (purple)).

Another highly relevant finding of our study is that the Hox TF Ubx lineage-specifically repressed the transcription of alternative fate genes by organizing the epigenetic landscape and that chromatin changes at Ubx sites were dependent on the interaction of Ubx with the Polycomb recruiter Pho. Strikingly, our study revealed that Ubx was crucial for stabilizing binding of the PcG protein Pho to specific chromatin regions. Although it is known that DNA binding TFs other than Pho interact with PREs, for example Grainyhead^74,75^, a TF that has been shown recently to lineage-specifically displace nucleosomes at enhancers^76^, these studies were mostly performed *in vitro* and the role of these TFs in Pho chromatin targeting was not addressed. In contrast, we analysed the combinatorial interaction of Ubx and Pho on the chromatin *in vivo*, in the mesodermal lineage of *Drosophila* embryos. Importantly, we showed that Pho was no longer able to interact with Ubx bound sites when Ubx protein was depleted in a lineage-specific manner, while Pho-only chromatin interactions were unaffected. As a result, histone modifications changed towards higher H3K27ac/H3K27me3 ratios, resulting in the de-repression of genes primarily associated with Ubx-Pho chromatin regions. This result indicated that not only Pho but very likely the whole PcG protein complex was not properly targeted to Ubx chromatin sites, which is in line with our finding that H3K27me3 levels were generally decreased at enhancers of genes de-repressed upon Ubx depletion (Fig. 7). However, we also found H3K27ac levels increased at Ubx chromatin interactions located in the vicinity of de-repressed genes. Interestingly, PREs frequently co-localize with response elements for Trithorax group (TrxG) proteins^52^, a large group of proteins first described for their role in transcriptional activation. In this line, we identified not only the binding motif for Pho but also for the GAGA TF Trithorax-like (Trl) enriched among the Ubx bound and H3K27me3 marked chromatin sites, and we found Trl to interact with Ubx *in cellulo* (Supplementary Fig. 5D). Trl was initially discovered to activate transcription, in particular of the *Ubx* gene^77^, but is also required for transcriptional repression, as it binds PREs^78^ and physically associates with the Polycomb Repressive Complex 1^79^. Interestingly, Trl had been suggested recently to function as a pioneer factor in early *Drosophila* development by making genomic regions accessible through the deposition of active histone marks. Furthermore, it had been shown that another TrxG protein, the methyltransferase Trithorax (Trx), mediates together with the p300/CREB-binding protein (CBP) the deposition of H3K27ac marks^80,81^, a histone mark that was enriched at promoters and enhancers upon Ubx depletion. Thus, the TrxG complex and CBP could be recruited in the absence of Ubx to activate gene expression (Fig. 7). GAGA factors like Trl do not only play a role in the activation but is also in the repression of gene expression by interacting with the Polycomb complex ^78,82^. Interestingly, a recent study showed that GAGA factors are required for the formation of repressive chromatin loops in Polycomb domains to stabilize gene silencing during early *Drosophila* development ^83^. These results could explain one puzzling result of our study, the increase of H3K27me3 marks in the absence of Ubx, which could be due to Trl interacting with these sites when Ubx levels drop (Fig. 7), thereby promoting or increasing the formation of repressive chromatin loops. In future, it will be important to study TrxG as well as PcG proteins at Ubx targeted control regions to understand how Hox TFs orchestrate the interplay between transcriptional activation and repression in the course of lineage development when cellular plasticity needs to be restricted.

One peculiar feature of Hox TFs is that they are not only active during the development but their input is continuously required throughout the lifetime of an organism to assess the positional values of cells and to maintain their proper identities and functions^84–90^. Our findings now shed new light on the mechanism ensuring this stability in cellular identity, as they suggest that Hox TFs robustly repress alternative lineage programs (via the interaction with epigenetic factors like Pho) and reliably restrict the plasticity also of adult cells. In the normal context, this is absolutely critical for an organism to function properly, however, in some instances it can be a major hurdle, in particular when cells need or should adopt a new identity, which requires them often to regain plasticity. In organismal life, this happens mostly during dedifferentiation, regeneration and tissue remodelling^91–93^. Another situation of cell type conversion is cellular reprogramming, which is extensively studied due to its high potential as therapeutic strategy^94^. But although reprogramming strategies have been improved over time, the direct conversion of one somatic cell type into another one, the so-called transdifferentiation, is still inefficient^6,95^. Interestingly, it has been reported recently that the induction of the Hox code typical for a differentiated cell type in pluripotent stem cells (PSCs), either in combination with other factors^96^ or even alone^11^, can convert PSCs into the cell type expressing the Hox code *in vivo*. This clearly shows that *Hox* genes can effectively induce cell type specific transcriptional programs in multi-potent cells thereby unambiguously inducing their differentiation. This is in line with our findings showing that Hox TFs activate lineage-specific transcriptional profiles with high precision. However, so far it is unclear why the conversion of somatic cells of one type into another one is so difficult. Our data now suggest that the main reason for this hurdle could be the Hox encoded restriction of cellular plasticity via the repression of alternative gene programs using Polycomb based epigenetic mechanisms.

Interestingly, it has been shown recently that the physiological conversion of intestinal to neuronal cells in *C. elegans* requires the removal of H3K27me3 marks to activate the motor neuron transcriptional programme^97^, a histone modification we find decreased at enhancer regions in the absence of Ubx. Thus, we postulate that effective cell type conversion in normal and experimental settings might require first the erasure of the Hox code established in the cells during development to silence the original gene program and to unleash the full potential of the cell by releasing repression of alternative programs. This would be followed by the enforced expression of a different Hox code, which will engage a new cell type specific identity program while repressing all other cell fates. And as *Hox* genes auto-regulatory control their own expression^98–102^, the Hox code should be maintained in the converted cells once initiated. Future experiments have to show whether this holds true and importantly whether this approach will enable us to effectively convert differentiated cell types *in vivo*.

## MATERIALS AND METHODS

### Fly stocks and husbandry

For the INTACT method^23^ the following fly lines were used: *twi-INTACT* and *UAS-INTACT* (gift from the Henikoff lab). The *UAS-INTACT* was crossed in the *elav-GAL4* background to generate *elav-GAL4;;UAS-INTACT* stable lines. For the degrade GFP experiments^20^ the following lines were used: *arm-GAL4* (BL1561) *Mef2-GAL4* (BL50742), *UAS-TEV-P14* (homemade using the construct published in Taxis et al., 2009, insertions were performed by BestGene at attP5 and attP2). *UAS-Nslmb-vhhGFP4* (BL38419), the *GFP-Ubx^3.005^* line was recombined with the *UAS-Nslmb-vhhGFP4* and *Mef2-GAL4* lines and crossed in the *twi-INTACT* background to generate *twi-INTACT;Mef2-GAL4,GFP-Ubx*^3.005^ and *twi-INTACT; UAS-Nslmb-vhhGFP4,GFP-Ubx^3.005^*. Embryos from Ubx deGradFP experiments were collected at 10-18h AEL to ensure the knockdown of the GFP-Ubx protein. The *pho-like*^*81A*^;*pho*^*1*^ douple mutant was generously provided by Jürg Müller^103^. The *dpp-lacZ* line was obtained from Manfred Frasch.

### Generation of endogenously tagged Ubx

The *GFP-Ubx*^3.005^ line was generating by using the CRIPR/Cas9 system^45,46^. For the donor DNA a pUC-MCS-5’GFP-MCS was designed, multiple cloning sites flanking the GFP containing an ATP start codon. For the 5’ region from the GFP a homologous arm^104^ containing the 5’UTR was cloned by using Not*I* and Kpn*I* restriction sites. The 3’ region from the GFP included the first Ubx exon and a large part of the intron for homologous recombination was cloned with Bgl*II* and Xho*I*. The gRNAs were designed to eliminate the first Ubx exon, positioned at the beginning of the 5’UTR and the end of the coding region of the first exon. The excised exon was replaced using the donor DNA and homologous recombination. The microinjection was performed by BestGene using *vas-Cas9* (BL51323) as injection line and the resulting progenies (F0) were crossed with TM3/TM6 balancers and resulting F1 was used for single crosses against TM3/TM6 balancers to generate independent stocks. The F1 generation was screened by PCR for the presence of the GFP and the GFP containing stocks of the F2 generation were visually screened for the Ubx patterned GFP expression *in vivo*.

### Generation of the Ubx antibody

The Ubx antibody was generated using the pGEX-purification system (GElifesciences). The open reading of Ubx-RA was cloned in the pGEX-6P-2 vector using Bam*HI* and Xho*I* restriction site. The protein was purified according to the protocol (GElifesciences) and eluted by using the PreScission Protease site. The immunisation and antibody handling was performed by the Charles Rivers company.

### Purification of affinity-tagged nuclei, ChIP, ChIP-Seq and RNA-Seq

The nuclei were purified as described in Steiner et al.^23^, the purification was optimised by using a larger magnet (20cmx20cmx10cm, holding 70kg) and a homemade magnet holder (gift from C. Schaub & M. Frasch). For ChIP and ChIP-Seq experiments: Chromatin preparation and immune-precipitation were performed as described in Sandmann et al., 2007^43^. The following antibodies were used: H3K27ac (ab4729, Abcam), H3K27me3 (ab6002, Abcam), Rb-Pho^2-382^ and the homemade guanine pig Ubx antibody (gp-Ubx). Total RNA was isolated with TRIZol (Invitrogen) and DNA digest was performed with the TURBO DNA-free Kit (Ambion). The material was analysis or validated by qPCR (Invitrogen Syber-Green-Mix) (Supplementary Tab. 2, 3). Genome-wide sequencing and material handling was performed in tight cooperation with the Deep-Sequencing facility in Heidelberg. The library for genome-wide sequencing was prepared by using the ThruPLEX DNA-Seq Kit (Rubicon) for illumina sequencing.

### Bioinformatics analysis

ChIP-Seq: The reads were first analysed with FastQC and aligned with bowtie2 (bowtie2 --very-sensitive -x Bowtie2Index/genome sequence.txt -S output.sam^105^ against the dm6 Drosophila genome version using standard conditions, the peaks were called by using MACS2 under model based broad settings (macs2 callpeak -t IP-file.sam -c input-file.sam -f SAM -n --broad -g dm --keep-dup auto -B --broad-cutoff 0.1 3. Further analysis and annotation of wildtype ChIP-Seq peaks were perfomed with ChIPseeker^106^, ChIPpeakAnno^107^. Annotations were not filtered for “real” coding genes and all FBgnNumbers were used for further analysis. Identification of differential Histone Binding events were identified with DiffBind, statistical relevant regions were isolated with BEDtools (bedtools intersect^108^). For GO-Term annotations and overrepresented GO-Term analysis was performed with the web-tools bioDBnet and PANTHER (GO biological function complete, Binomial, Bonferroi correction^109^. Promoter and enhancer definition is related to ^15^ and ^64^.

Motif search: The motif search on defined regions was done by using the MEME-suite web-tool^110^ or the command line (meme-chip-db motif_databases/FLY/OnTheFly_2014_Drosophila.meme -meme-minw 6 -meme-maxw 20 -meme-nmotifs 100 -centrimo-local -centrimo-maxreg 250 seq.fa). Enrichment of known motifs was performed with a AMT MEME tool (ame --verbose 1 --oc. --control shuffled_overlaps_onlyPho_ME_2.fa --bgformat 1 --scoring avg --method ranksum -- pvalue-report-threshold 0.05 file.fa db/FLY/fly_factor_survey.meme db/FLY/idmmpmm2009.meme db/FLY/flyreg.v2.meme db/FLY/OnTheFly_2014_Drosophila.meme db/FLY/dmmpmm2009.meme) Comparison of Ubx and Tin: The Tin data set from Jin et al., 2013 was used, for that purpose the Ubx peaks were realigned and called against Drosophila genome dm3. Analysis of Ubx and Pho ChIP-Seq data: The Pho data set was obtained from Erceg et al., 2017 and processed as decried above for Ubx. Genomic overlaps were performed with ChIPpeakAnno^107^ using a maxgap of 250 bp.

RNA-Seq: Obtained RNA reads were first analysed with FastQC and aligned with TopHat2^111^ (tophat2 -p 8 -G Drosophila_melanogaster.BDGP6.86.gtf -o output_folder_name BDGP6_86 converted_sequence.fastq) against the BDGP6_86 Drosophila genome version using default settings. The count table for further analysis was generated using HTSeq (htseq-count -f bam -r pos -m union -s no -t exon accepted_hits.bam Drosophila_melanogaster.BDGP6.86.gtf > counts.txt ^112^. The differential expression analysis was performed by using DESeq2^113^. The overrepresented GO-Term were analysed with PANTHER (GO biological function complete, Binomial, Bonferroi correction).

### Visualisation of the genome-wide data

Preparation of the data for visualisation was perfomed with deeptools: Normalization computeGCBias -b file_sorted.bam --effectiveGenomeSize 142573017 -g dm6.2bit -l 250 --GCbiasFrequenciesFile file_freq.txt --biasPlot file_test_gc.pdf and correctGCBias -b file_sorted.bam --effectiveGenomeSize 142573017 -g dm6.2bit -- GCbiasFrequenciesFile file_freq.txt -o file_gc_corrected.bam. The samples were subtracted from the input to generate the bigwig file bamCompare -b1 sample_gc_corrected.bam -b2 input_gc_corrected.bam -o file_S-I_100_subtract.bw -- scaleFactorsMethod None --operation subtract --binSize 10 --effectiveGenomeSize 142573017 --normalizeUsing RPKM --smoothLength 100 --centerReads -- extendReads 150. The reads and annotations were visualised with IGV.

### Interactive data mining tool

The enrichment analysis method presented in this paper is implemented as a user-friendly Shiny web-application accessible via http://beta-weade.cos.uni-heidelberg.de.. The user can select the set of genes to perform the GO enrichment analysis and the respective background independently. Results of the analysis are presented as a plot, an interactive table displaying significantly enriched GO groups, and an interactive heatmap, showing the counts of enriched GO terms within the respective higher-order GO group. It is also possible to get an insight into the individual GO terms that make up a category and into the genes that contributed to the categories or terms. The functionality of the tool exceeds what is described here, a detailed documentation of the tool is deposited under http://beta-weade.cos.uni-heidelberg.de. and additionally an interactive guide is provided in the online application.

#### Multiple testing of GO-terms

Pre-set list of GO-terms related to mesoderm and muscle development and function (mesoderm), neuronal and central nervous system (CNS) and neuromuscular junction development and function (neuronal), ectoderm and trachea development and function (ectoderm) and endoderm development and gut structures (salivary glands, Malpighian tubes, hind-and foregut) and function (gut & endoderm) was generated (Supplementary Tab. 4). Genes associated with different categories were analysed using an R program based on a biomaRt script^114^.

### Immunofluorescence staining and microscopy

Nuclei staining: INTACT-sorted nuclei were collected by centrifugation at 1000 g for 5 min and resuspend in the suitable amount of HB125 (Steiner et al.^23^) +5 % BSA for the antibody staining according to the conditions of the 1^st^ antibodies and incubated over night at 4 °C, than washed 3 times 15 min with HB125 and incubated in 500 µl HB125 +5 % BSA for the 2^nd^ antibodies (2^nd^ ABs 1:500). Before mounting, nuclei were washed 3 times 15 min with HB125. Embryonic stainings were performed as described in Domsch et al., 2013^115^ and the following antibodies were used: Rb-Mef2 (1:1500, Gift from H. Nguyen), Rat-elav (1:50, DSHB, ELAV-9F8A9), Rat-Tm1 (1:200, MAC-141, ab50567, Abcam), Rb-GFP (1:300, Invitrogen), Rat-GFP (1:100, ChromoTek), DAPI (1:500, Invitrogen), Rat-Org1 (1:100)^92^, Rb-Pho (1:250)^103^, Ptx1 antibody (1:1000)^69^, ms-Antp (1:100, DSHB, 8C11). Amplification was obtained with the TSA system (Perkin-Elmer). Biotinylated (1:200, Vector Labs) and fluorescent (1:200, Jackson ImmunoResearch) secondary antibodies were used. Images were acquired on the Leica SP9 Microscope using a standard 20× and 63× objective. The collected images were analysed and processed with the Leica program and Adobe Photoshop.

### Quantifications of data

Quantification of the GFP signal intensity: GFP signal was measured with the Fiji program from at least 5 segments per embryo in 5 embryos. The mean and the t-test for significance were calculated in Excel.

Quantification of histone marks: Regions of interest (Ubx bound regions with histone mark changes that are up-regulated or without up-regulation, Ubx and Pho bound regions, Ubx Pho and active regions) were identified and compared with the deGrad-DiffBind results, histone binding peaks from control or deGrad-data stets by using bedtools intersect or bedtools coverage. The isolated Fold or peak depth were illustrated in a Box-plot with R, as well as the wilcoxon rank test for significance. Fold changes required for ratio calculation were processed with an R-script (upon request) and illustrated in a Box-plot with R, as well as the Wilcoxon rank test for significance.

### Protein IP, CoIP and Western Blot

Embryonic Protein IP: The GFP-Ubx protein was immune precipitated from whole mount embryos by using GFP-tap-beads (ChromoTek). Collected Embryos were washed and smashed in IP Buffer (50 mM Tris-HCl, pH 8.0; 1 mM EDTA; 0.5% (v/v) Triton X-100, 1 mM PMSF, oComplete proteinase inhibitors (ROCHE). For immunoprecipitation 1 mg of total protein was incubated over night with IP Buffer washed GFP-tap-beads. Beads were than washed 3x with IP Buffer and resuspended in Laemmli buffer.

Cell culture CoIP: S2R+ Drosophila cells were cultured in Schneider medium supplemented with 10% FCS, 10U/ml penicillin and 10µg/ml streptomycin. For plasmid transfections, 10.10^6^ cells per ml were seeded in 100mm dishes and transfected with Effectene (Qiagen) according to the manufacturer’s protocol. Cells were harvested in Phosphate Buffered Saline (PBS) and pellets were resuspended in NP40 buffer (20mM Tris pH7.5, 150mM NaCl, 2mM EDTA, 1% NP40) supplemented with protease inhibitor cocktail (Sigma-Aldrich) and 1mM of DTT. For co-immunoprecipitation assays 1mg to 1.5mg of proteins were incubated for 3h with 15μl of GFP-Trap beads (Chromotek). Beads were then washed 3 times with NP40 buffer and finally resuspended in Laemmli buffer for immunoblotting analysis. Input fractions represent 2.5% of the immunoprecipitated fraction.

Embryonic CoIP: Overnight collection of embryos was dechorionated, fixed with 3.2% formaldehyde and collected in Phosphate Buffered Saline (PBS) supplemented with Tween 0.1%. Pellets were resuspended in buffer A (10 mM Hepes pH 7.9, 10 mM KCl, 1.5 mM MgCl2, 0.34 M sucrose, 10% glycerol) and dounced 25-30 times with loose-and 5 times with tight-pestle. Lysates were incubated with 0.1% Triton and centrifugated. Nuclear pellet were then resuspended with buffer B (3mM EDTA pH8, 0.2mM EGTA pH8), sonicated (Picoruptor, Diagenode), and treated with Benzonase (Sigma). For co-immunoprecipitation assays 4 to 5 mg of nuclear lysates were diluted in NP40 buffer (20 mM Tris pH7.5, 150 mM NaCl, 2 mM EDTA, 1% NP40) and incubated overnight with 40 µl of GFP-Trap beads (Chromotek). Beads were then washed 5 times with NP40 buffer and resuspended in Laemmli buffer for immunoblotting analysis. All buffers were supplemented with protease inhibitor cocktail (Sigma) and 1mM of DTT and 0.1mM PMSF. Input fractions represent 2.5% of the immunoprecipitated fraction.

For western blot analysis, proteins were resolved on 10% SDS-PAGE, blotted onto PVDF membrane (Biorad) and probed with specific antibodies after saturation. The antibodies (and their dilution) used in this study were: Ubx (home-made, 1/200), Pho (generously provided by Jürg Müller, 1/500 ^103^, Histone 3 (ab1791, Abcam, 1/10000), GFP (A-11122, Life Technologies, 1/3000).

## ACKNOWLEDGMENTS

We would like to thank the *Drosophila* community for providing us generously with fly stocks and antibodies, in particular Steven Henikoff for the *twi-INTACT* and *UAS-INTACT* fly lines, Jürg Müller for the Pho antibody and the *phol*^*81A*^;*pho^1^* double mutants and Gerd Vorbrüggen for the Ptx-1 antibody. Manfred Frasch and Christoph Schaub at the University of Erlangen-Nuremberg in Germany for the improved INTACT apparatus. David Ibberson from the Heidelberg Deep Sequencing Core facility at Heidelberg University and EMBL. Eugen Rempel for the Bioinformatics support. We also thank all the members of the lab for critically reading the manuscript. We apologize to all whose work was not cited due to space limitations. This work was supported by the DFG (DFG, LO 844/8-1).

## AUTHOR CONTRIBUTIONS

K.D. conceived, designed, performed and interpreted genome wide RNA-Seq, Ubx ChIP-Seq, histone ChIP-Seq as well as Ubx degradation RNA-Seq and ChIP-seq experiments. K.D. conceived, designed and isolated together with V.D. the endogenous GFP-Ubx line. J.C. performed histone ChIP-Seq and Co-IP experiments in cell culture and whole *Drosophila* embryo. M.P. generated the Ubx antibody used in this study. K.D and J.F performed and interpreted in situ experiments. N.T. and O.E. wrote bioinformatics scrips and helped with the statistical analysis of the genome wide data. I.L. conceived the study, assisted in designing and interpreting experiments, wrote the paper together with K.D. and obtained funding to support the study (DFG, LO 844/8-1).

